# PANDA: A comprehensive and flexible tool for proteomics data quantitative analysis

**DOI:** 10.1101/332957

**Authors:** Cheng Chang, Chaoping Guo, Yuqing Ding, Kaikun Xu, Mingfei Han, Fuchu He, Yunping Zhu

## Abstract

**Summary:** As the experiment techniques and strategies in quantitative proteomics are improving rapidly, the corresponding algorithms and tools for protein quantification with high accuracy and precision are continuously required to be proposed. Here, we present a comprehensive and flexible tool named PANDA for proteomics data quantification. PANDA, which supports both label-free and labeled quantifications, is compatible with existing peptide identification tools and pipelines with considerable flexibility. Compared with MaxQuant on two complex datasets, PANDA was proved to be more accurate and precise with less computation time. Additionally, PANDA is an easy-to-use desktop application tool with user-friendly interfaces.

**Availability:** PANDA is freely available for download at https://sourceforge.net/projects/panda-tools/.

## 1. INTRODUCTION

Quantitative proteomics is gaining its popularity by providing a global and systematic view on biological processes and cellular functions (Schubert, et al., 2017). There are two kinds of approaches to protein quantification according to whether the sample is isotope labeled, i.e. the label-free and labeled quantifications. Nowadays, numbers of algorithms and software tools have been proposed and developed to facilitate label-free or labeled quantification of proteomics data.

Due to the variety of experiment designs and strategies in quantitative proteomics, current quantification software tools are usually only suitable for a few specific quantitative experiment strategies, such as PyQuant (Mitchell, et al., 2016) and SILVER (Chang, et al., 2014) for stable isotope labeling quantification, RIPPER (Van Riper, et al., 2016) and LFQuant (Zhang, et al., 2012) for label-free quantification. Even the famous tool MaxQuant (Cox and Mann, 2008), which contains many methods for label-free and labeled quantifications, cannot support 15N labeling method. Moreover, MaxQuant consists of its own mass spectrometry (MS) data analysis algorithms, which are not compatible with other tools or pipelines. In brief, there is a lack of comprehensive and flexible quantification tools for the rapidly developing quantitative proteomics.

Here, we present a new tool named PANDA for accurate and precise analysis of quantitative proteomics with high comprehensiveness and flexibility. PANDA can process MS data from different instrument manufacturers by reading the standard formats mzXML and mzML. It is also able to be compatible with existing peptide identification tools (e.g. Mascot) and proteomics data analysis pipelines, such as the Trans-Proteomic Pipeline (Deutsch, et al., 2010). PANDA contains multiple methods to deal with MS data produced in various kinds of quantitative strategies. Further, by integrating the advanced algorithms of our previous quantification tools LFQuant and SILVER, PANDA has been demonstrated to be accurate and precise for protein quantification.

## 2. METHODS

### Benchmark datasets

For label-free quantification, the yeast samples with a serial dilution of UPS2 (Proteomics Dynamic Range Standard, SigmaAldrich) standard proteins (1µg, 0.2µg, 0.04µg, 0.008µg) spiked in from (Chang, et al., 2016) were analyzed in this study. For labeled quantification, a large-scale complex dataset obtained from HeLa cells (Cox and Mann, 2007) with stable isotope labeling by amino acids in cell culture (SILAC) was used. See Supplementary Methods for details.

### PANDA workflow

PANDA is designed for comprehensive and flexible analysis of both label-free and labeled quantitative proteomics data. As shown in Fig. 1, PANDA consists of three core layers, i.e. the data layer, the function layer and the algorithm layer. (1) The data layer includes two kinds of input data in PANDA: MS data and peptide identifications. For MS data, PANDA can directly process Thermo raw files through MSFileReader. Besides, it can also take the MS data standard formats mzXML and mzML as input. For peptide identification, being able to access the mzIdentML format proposed by the Human Proteome Organization Proteomics Standards Initiative makes it possible for PANDA to quantify the results of the commonly-used peptide identification tools, such as Mascot, SEQUEST, X!Tandem and MS-GF+. Meanwhile, PANDA can read the quality control results of PeptideProphet (Keller, et al., 2002) and PepDistiller (Li, et al., 2012), which further broadens its usage and flexibility. (2) The function layer contains the current mainstream quantification methods. For label-free quantification, spectral count (SC) method and extracted-ion chromatography (XIC) (also named as intensity-based) method were implemented in PANDA. As to labeled quantification, PANDA supports the prevalent precursor ion labeling methods, i.e., SILAC, 18O, 15N, isotope-coded affinity tags (ICAT) and isotope-coded protein labels (ICPL), as well as product ion labeling methods, i.e. isobaric tag for relative and absolute quantitation (iTRAQ) and tandem mass tag (TMT). Furthermore, users can define their own labeling methods in PANDA. (3) The algorithm layer includes the basic algorithms for MS data processing and peptide/protein quantification. Part of them are adapted from LFQuant and SILVER, such as the reversible retention time (RT) alignment algorithm in LFQuant, the multi-filters for XIC construction and the dynamic isotopic matching tolerance algorithm in SILVER.

**Fig. 1.**
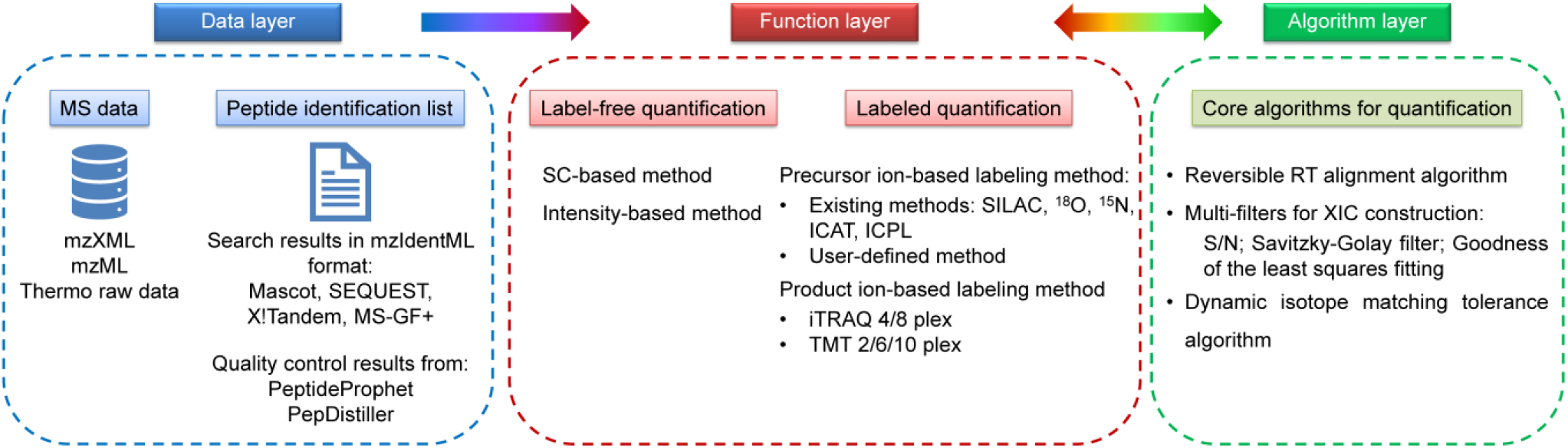
The schema of PANDA workflow.

## 3 RESULTS

In this study, PANDA was compared with MaxQuant (v1.6.0.13, released on Aug 2017) on a yeast dataset with four concentration levels of UPS2 standard proteins spiked in (A-D groups) and a large-scale HeLa dataset with SILAC labeling for label-free and labeled quantifications, respectively.

### Accuracy evaluation

In the yeast dataset, the theoretical ratios of the spiked-in UPS2 proteins for A/B, A/C and A/D should be 5, 25, 125. As shown in Supplementary Figure 1, the quantification results of PANDA were closer to the theoretical ratios than those of MaxQuant. In the HeLa dataset, the SILAC ratios of the 3471 proteins commonly quantified by PANDA and MaxQuant were shown in Supplementary Figure 2. The ratio distribution of PANDA was also closer to the theoretical ratio (1:1) than that of MaxQuant. These results demonstrated that PANDA has a high accuracy for both label-free and labeled quantifications in a wide dynamic range.

### Precision evaluation

In the yeast dataset, PANDA showed a lower coefficient of variation (CV) distribution of the yeast proteins for the technical replicates within each group (A-D) than MaxQuant, indicating the high precision of PANDA for label-free quantification (Supplementary Figure 3). In the HeLa dataset, the protein intensity CVs of the three technical replicates for both SILAC labeled and unlabeled samples were calculated and PANDA also displayed a lower CV distribution than MaxQuant, which proved that PANDA is precise for labeled quantification (Supplementary Figure 4). More details are provided in Supplementary Notes.

Finally, PANDA is efficient due to the refinement of its source codes and the inclusion of popular third-party libraries, such as GNU scientific library. It spent less computation time than MaxQuant on the two datasets (Supplementary Table 1).

## 4 CONCLUSION

In summary, PANDA contains a comprehensive algorithm collection for label-free and labeled quantifications and supports all the main methods in quantitative proteomics. Being able to read proteomics data in public format, PANDA is very flexible and compatible with existing peptide identification tools or MS data analysis pipelines. Most importantly, PANDA is proved to be accurate and precise for label-free and labeled quantifications. At last, the quantification results of PANDA can be further analyzed in its affiliated tool PANDA-view (Chang, et al., 2018) for statistical analysis and data visualization.

## Funding

This work was supported by the National Key Research and Development Program of China [2017YFA0505002 and 2017YFC0906602] and the National Natural Science Foundation of China [21605159 and 21475150].

## Conflict of Interest

none declared.

